# CSF glucosylsphingosine is a central readout of GCase impairment across genetic and sporadic Parkinson’s disease

**DOI:** 10.64898/2026.06.19.733447

**Authors:** Jillian H. Kluss, Di Jiang, Shan V. Andrews, Ritesh Ravi, Romeo Maciuca, Buyankhishig Tsogtbaatar, Chau Tran, Meng Fang, Eric A. Macklin, Anastasia G. Henry, Michael A. Schwarzschild, Sarah Huntwork-Rodriguez

## Abstract

Parkinson’s disease (PD) risk converges on lysosomal biology, including reduced activity of glucocerebrosidase (GCase), encoded by *GBA1*. Glucosylsphingosine (GlcSph), a toxic deacylated glycosphingolipid and established target engagement biomarker in Gaucher disease, is difficult to quantify in cerebrospinal fluid (CSF) because of low abundance and isomeric interference. We optimized and qualified targeted LC–MS/MS assays for GlcSph in CSF and plasma and measured GlcSph and GCase activity (4-MU assay) in participants from the Parkinson’s Progression Markers Initiative (PPMI), including *GBA1* and *LRRK2* mutation carriers with and without PD, sporadic PD, and healthy controls. Group level analyses showed higher levels of CSF and plasma GlcSph in *GBA1* heterozygous variant carriers independent of disease status, ∼120% (p<0.0001) and ∼61% (p<0.0001), respectively, with CSF elevations reflecting allelic dosage and, to a lesser degree, variant severity. Notably, elevated CSF GlcSph was observed not only in carriers of pathogenic *GBA1* mutations but also in individuals harboring common *GBA1* risk variants. CSF GlcSph was also elevated by ∼30% in sporadic PD (p=0.002) and by ∼40% *LRRK2* mutation carriers (p=0.0003), indicating shared central GCase pathway perturbation across PD subtypes. CSF and plasma GlcSph levels showed poor concordance across groups with the exception of the GBA-PD group, supporting the hypothesis that central and peripheral GCase pathway dysfunction arise through distinct biological mechanisms. Together, these findings establish CSF GlcSph as a sensitive biomarker of central GCase pathway impairment and may support its use as a pharmacodynamic and target engagement biomarker for therapeutics targeting GCase in PD. More broadly, CSF GlcSph may enable identification of biologically defined GCase-pathway dysfunction beyond genetically defined *GBA1*-associated PD, with potential implications for patient stratification in future disease-modifying trials.

## Introduction

Parkinson’s disease (PD) is a common, progressive neurodegenerative disease with clinical and biological variation^1^. This is most likely the result of compounding effects of an individual’s age, environmental exposures, and genetic profile. Despite the heterogeneity in the manifestations of PD, genetic and biological evidence strongly supports the convergence of many risk factors into a few key biological pathways. One such pathway is the endolysosomal pathway, where genome-wide association studies (GWAS) have revealed variants in several lysosomal enzymes and associated transporters that are significantly associated with disease risk^2–4^. *GBA1* (referred to as GBA for the remainder of this manuscript) is one of these genes, in which heterozygous mutations result in reduced enzyme activity of the lysosomal protein glucocerebrosidase (GCase). Consistent with this genetic evidence, several studies have shown reduced GCase protein and enzymatic activity in the brains of individuals with sporadic PD compared to healthy controls^5–8^. Taken together, the available evidence supports a shared signature of lysosomal perturbation present across various manifestations of PD, with GCase likely playing a central role.

Interestingly, autosomal recessive mutations in GBA cause Gaucher Disease (GD), a lysosomal storage disorder hallmarked by organomegaly, bone fragility, and blood cell abnormalities. These loss of function mutations lead to the accumulation of several glycosphingolipid species including a toxic lipid substrate glucosylsphingosine (GlcSph) that is formed from the deacylation of glucosylceramide (GlcCer), the primary substrate of GCase^9^. GlcSph accumulates in the peripheral tissues and fluids of individuals with GD, as well as in the brain tissue of those with neuronopathic forms of the disease^10,11^. In addition, several studies have shown elevated levels of GlcSph in plasma and red blood cells in individuals with GD compared to controls, some of which reported differences greater than 200-fold magnitude^12–15^. In clinical studies, restoration of GCase activity via enzyme replacement therapy (ERT) ameliorates GlcSph levels to within reference range, establishing plasma GlcSph as a direct readout of GCase activity and increasing its utility as a target engagement biomarker for the diagnosis and monitoring of GD^13,15,16^.

In PD, one published study has reported elevated levels of plasma GlcSph in GBA mutation carriers compared to non-carriers, suggesting that GlcSph may have utility as a biomarker in GBA-PD^17^. In contrast to plasma, measurement of brain-derived GlcSph in CSF has proven challenging, as levels of GlcSph are naturally low in cerebrospinal fluid (CSF) while levels of galactosylsphingosine (GalSph), the structural isomer of GlcSph, are significantly higher^18^, requiring a sensitive and selective assay. To fill this gap and evaluate the utility of GlcSph as a biomarker of PD, we optimized and qualified two targeted liquid chromatography with tandem mass spectrometry (LC-MS/MS) assays^18^ for detection of GlcSph in CSF and plasma, respectively. Using cohorts of 568 CSF and 640 plasma samples from the Parkinson’s Progressive Markers Initiative (PPMI), we measured GlcSph levels in *GBA* and *LRRK2* carriers with and without PD as well as sporadic cases and healthy controls. Additionally, we measured CSF and plasma GCase activity via the 4-methylumbelliferyl-B-D-glucoside (4-MU) assay in 640 CSF and plasma samples and correlated all four analytes with available demographic and selected clinical assessment data to understand the scope of utility of these biomarkers.

## Results

### CSF GlcSph is elevated in sporadic PD and monogenic carriers

To assess the levels of GlcSph in human CSF and plasma utilizing the optimized LC-MS/MS targeted assays, we analyzed 568 CSF samples (collected from 98 healthy controls, 98 sPD, 111 LRRK2-NMC (non-manifesting carrier), 110 LRRK2-PD, 91 GBA-NMC, 60 GBA-PD) and 640 plasma samples (collected from 113 healthy controls, 106 sPD, 115 LRRK2-NMC, 117 LRRK2-PD, 87 GBA-NMC, 91 GBA-PD, 7 LRRK2/GBA-NMC, 4 LRRK2/GBA-PD) from PPMI participants. These samples were selected for balanced age, sex, and disease duration at the time of sample collection across all groups, and for availability of paired CSF and plasma samples where feasible (additional details on demographics can be found in Tables 1 and 2). Within each LRRK2 and GBA genetic group, all variants were heterozygous except for 11 homozygous N370S GBA carriers. There were also a few individuals that were carriers of a GBA common risk variant such as E326K or T369M in addition to a G2019S LRRK2 mutation (5 in the CSF group and 8 in the plasma group). For simplicity, these individuals were treated as LRRK2 carriers only. There were also 11 carriers harboring both G2019S LRRK2 and N370S GBA pathogenic mutations in the plasma cohort which were excluded from the group level analyses. For the performance characteristics of both quantitative assays, refer to Table 3.

**Table 1.** CSF cohort demographics. Average disease duration and Median age of sample median are reported in years. HC: Healthy Control; sPD: sporadic Parkinson’s Disease; LRRK2-NMC: LRRK2-positive non-manifesting carrier; LRRK2-PD: LRRK2-positive Parkinson’s Disease; GBA-NMC: GBA-positive non-manifesting carrier; GBA-positive Parkinson’s Disease.

|  | **HC**<br>(N=116) | **sPD**<br>(N=110) | **LRRK2-NMC**<br>(N=120) | **LRRK2-PD**<br>(N=120) | **GBA-NMC**<br>(N=104) | **GBA-PD**<br>(N=70) | **Total Subjects**<br>(N=640) |
| --- | --- | --- | --- | --- | --- | --- | --- |
| *Female* | 51 (44.0%) | 47 (42.7%) | 55 (45.8%) | 55 (45.8%) | 48 (46.2%) | 28 (40.0%) | 284 (44.4%) |
| *Male* | 65 (56.0%) | 63 (57.3%) | 65 (54.2%) | 65 (54.2%) | 56 (53.8%) | 42 (60.0%) | 356 (55.6%) |
| *Average age*<br>*(SD)* | 62.3 (9.9) | 62.4 (8.5) | 62.6 (7.7) | 63.2 (8.4) | 62.1 (6.9) | 61.1 (9.6) | 62.4 (8.5) |
| *Average*<br>*disease*<br>*duration (SD)* | 0 (0) | 2.87 (2.12) | 0 (0) | 2.96 (2.17) | 0 (0) | 3.08 (2.37) | 2.96 (2.2) |
| *Matching*<br>*plasma GCase*<br>*activity sample* | 74 (72.5%) | 88 (89.2%) | 96 (86.7%) | 98 (68.3%) | 54 (61%) | 42 (41.7%) | 457 (72.8%) |
| *CSF GlcSph*<br>*sample* | 98 (84.5%) | 98 (89.1%) | 111 (92.5%) | 110 (91.7%) | 91 (87.5%) | 60 (85.7%) | 568 (88.8%) |
| *Matching*<br>*plasma GlcSph*<br>*sample* | 69 (59.5%) | 87 (79.1%) | 95 (79.2%) | 77 (64.2%) | 54 (51.9%) | 26 (37.1%) | 408 (63.8%) |
| *Median age of*<br>*sample*<br>*(Q1, Q3)* | 11.1<br>(9.9, 12.2) | 9.60<br>(8.12, 11.0) | 6.50<br>(5.60, 7.43) | 8.20<br>(6.50, 9.10) | 5.10<br>(4.30, 6.40) | 7.45<br>(6.32, 8.17) | 7.70<br>(6.10, 10.0) |

**Table 2.** Plasma cohort demographics. Average disease duration and Median age of sample are reported in years. HC: Healthy Control; sPD: sporadic Parkinson’s Disease; LRRK2-NMC: LRRK2-positive non-manifesting carrier; LRRK2-PD: LRRK2-positive Parkinson’s Disease; GBA-NMC: GBA-positive non-manifesting carrier; GBA-positive Parkinson’s Disease.

|  | **HC**<br>(N=113) | **sPD**<br>(N=106) | **LRRK2-**<br>**NMC**<br>(N=115) | **LRRK2-**<br>**PD**<br>(N=117) | **GBA-NMC**<br>(N=87) | **GBA-PD**<br>(N=91) | **LRRK2/**<br>**GBA-NMC**<br>(N=7) | **LRRK2/**<br>**GBA-PD**<br>(N=4) | **Total**<br>**Subjects**<br>(N=640) |
| --- | --- | --- | --- | --- | --- | --- | --- | --- | --- |
| *Female* | 48<br>(42.5%) | 47<br>(44.3%) | 61<br>(53.0%) | 60 (51.3%) | 44 (50.6%) | 40 (44.0%) | 2<br>(28.6%) | 3<br>(75.0%) | 305<br>(47.7%) |
| *Male* | 65<br>(57.5%) | 59<br>(55.7%) | 54<br>(47.0%) | 57 (48.7%) | 43 (49.4%) | 51 (56.0%) | 5<br>(71.4%) | 1<br>(25.0%) | 335<br>(52.3%) |
| *Average age*<br>*(SD)* | 63.9 (9.5) | 64.3 (8.4) | 64.6 (7.7) | 64.3 (8.5) | 63.9 (8.5) | 63.8 (8.9) | 69.4 (9.2) | 58.8 (11) | 64.2 (8.6) |
| *Average*<br>*disease*<br>*duration*<br>*(SD)* | 0 (0) | 2.77<br>(2.06) | 0 (0) | 3.30 (2.29) | 0 (0) | 3.71 (2.21) | 0 (0) | 3.84 (2.22) | 3.25 (2.22) |
| *Median age*<br>*of sample*<br>*(Q1, Q3)* | 8.77<br>(7.86,<br>9.96) | 7.54<br>(6.04,<br>8.51) | 4.27<br>(3.63, 5.23) | 4.83<br>(3.85,<br>6.41) | 3.03<br>(2.52,<br>4.26) | 4.11<br>(3.11,<br>5.58) | 3.96<br>(2.04,<br>4.21) | 5.83<br>(4.89,<br>6.76) | 5.36<br>(3.84,<br>7.68) |

**Table 3.** CSF and Plasma Glucosylsphingosine assay qualification specifications. The intra- and inter-assay CVs were calculated as percent recovery from duplicate measures of a series of QCs of known concentrations. Long-term stability of these assays is currently being established.

|  | CSF assay qualification | Plasma assay qualification | CSF PPMI | Plasma PPMI |
| --- | --- | --- | --- | --- |
| *Qualified dynamic range* | 0.1 - 50 pg/mL | 0.1 - 10 ng/mL | -- | -- |
| *Intra-assay CVs* | 1.9 - 13.6% | 1.6 - 6.3% | 0 – 27% | 0.3 - 4.9% |
| *Inter-assay CVs* | 3.8 - 10.2% | 2.8 - 3.4% | 2 – 8%<br>(across 10 plates) | 3.1%<br>(across 22 plates) |
| *Freeze/thaw stability* | Stable up to 4<br>freeze/thaws | Stable up to 5 freeze/thaws | -- | -- |
| *Benchtop stability* | ≤ 19 hours at RT | ≤ 24 hours at RT | -- | -- |

To understand the effect of GBA mutations on CSF GlcSph without disease influence, we compared levels between GBA-NMCs and healthy controls. We observed a 141% (p<0.0001) increase in the GBA-NMC group compared to healthy controls calculated from ANCOVA pairwise tests adjusted for age, sex, and batch run with Benjamini–Hochberg correction for multiple comparisons (Fig. 1A). When omitting the N370S homozygous carriers from the GBA-NMC group, this increase was reduced to 122% (p<0.0001). Plasma GlcSph levels were increased by 69% (p<0.0001) in GBA-NMCs compared to healthy controls, which was reduced to 61% when omitting the N370S homozygous carriers (Fig. 1B). CSF GlcSph was also increased in both sporadic PD and LRRK2-NMC groups by 32% (p=0.002) and 37% (p=0.0003), respectively, compared to healthy controls. These statistical differences were not observed in the plasma, as sPD and LRRK2 carrier groups exhibited similar GlcSph levels to healthy controls. (Fig. 1B). Within the GBA and LRRK2 carrier groups, there was no significant difference between NMC and PD groups, suggesting that disease status does not influence GlcSph levels, in either CSF or in plasma (Fig. 1A, B). A summary of all unadjusted absolute values can be found in Table 4.

**Figure 1.**
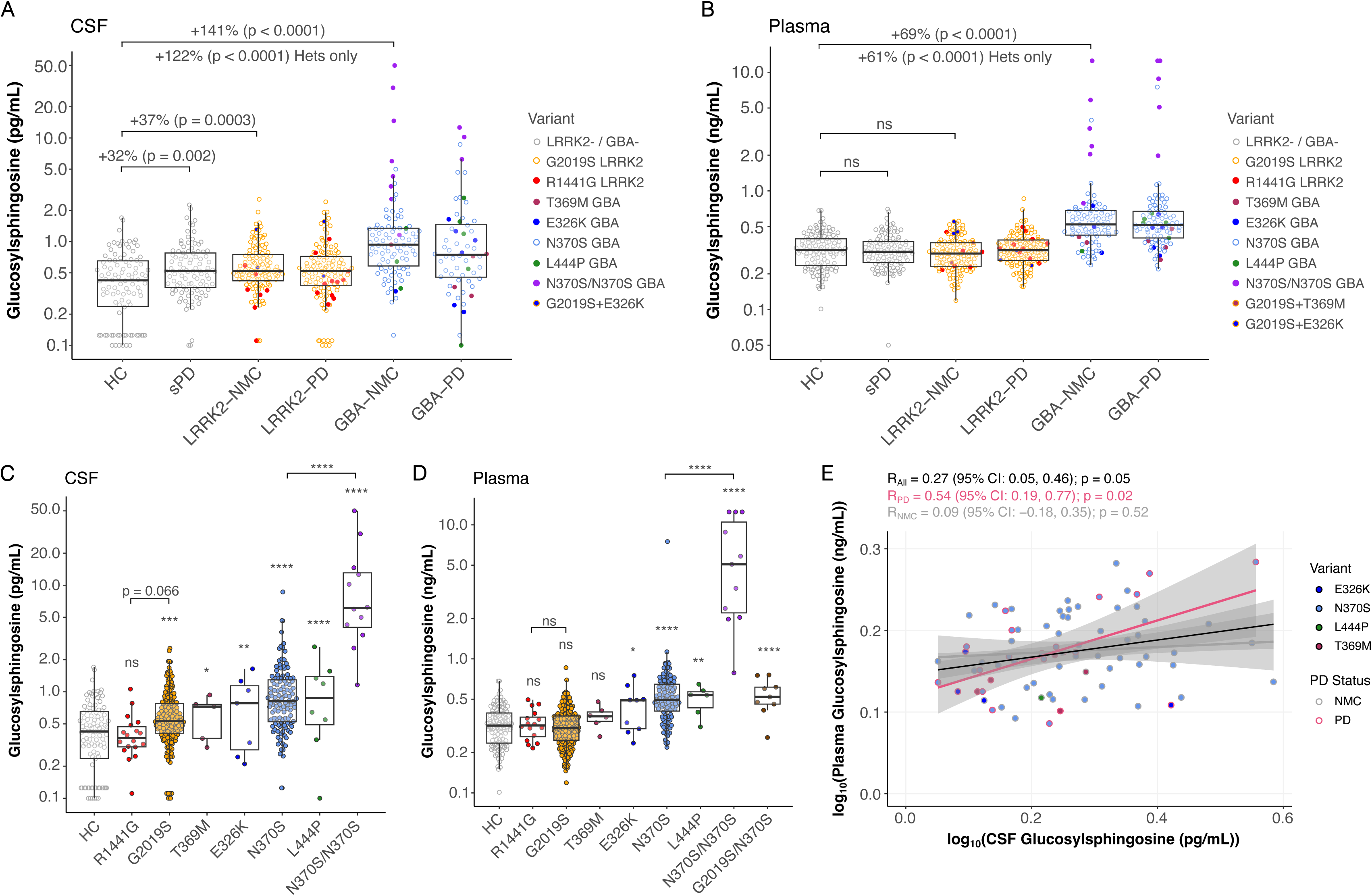
CSF and plasma GlcSph are elevated in GBA mutation carriers and selectively increased in CSF in sporadic PD and LRRK2 G2019S carriers. **A** CSF and **B** plasma GlcSph concentrations measured by targeted LC–MS/MS across healthy controls (HC), sporadic Parkinson’s disease (sPD), LRRK2 non-manifesting carriers (LRRK2-NMC), LRRK2-PD, GBA non-manifesting carriers (GBA-NMC), and GBA-PD. **C** CSF and **D** plasma GlcSph are stratified by individual genetic variants. The asterisks above each boxplot are in relation to HCs if not specified with a bracket. **E** Correlation between paired CSF and plasma GlcSph levels within GBA carriers. Data are shown as boxplots with individual data points overlaid; boxes represent the interquartile range with median indicated, and whiskers denote 1.5× IQR. Statistical comparisons were performed using ANCOVA pairwise tests with Benjamini–Hochberg correction for multiple comparisons; adjusted *p*-values are shown. Correlations were assessed using Pearson correlation on log-transformed values. Asterisks denote significance; * p < 0.05, ** p < 0.01, *** p < 0.001.

**Table 4.** Descriptive statistics derived from the unadjusted absolute values of the CSF and plasma GlcSph cohorts.

| Subgroup | Disease Status | Variant | N<sub>CSF</sub> | Median<sub>CSF</sub>(pg/mL) | Mean<sub>CSF</sub>(pg/mL) | Geometric Mean<sub>CSF</sub> (pg/mL) | SD<sub>CSF</sub> (pg/mL) | N<sub>Plasma</sub> | Median<sub>plasma</sub> (ng/mL) | Mean<sub>plasma</sub> (ng/mL) | Geometric Mean<sub>plasma</sub> (ng/mL) | SD<sub>Plasma</sub> (ng/mL) |
| --- | --- | --- | --- | --- | --- | --- | --- | --- | --- | --- | --- | --- |
| HC | - | - | 98 | 0.42 | 0.48 | 0.37 | 0.33 | 113 | 0.32 | 0.33 | 0.31 | 0.11 |
| sPD | PD | - | 98 | 0.52 | 0.62 | 0.52 | 0.39 | 106 | 0.31 | 0.33 | 0.31 | 0.11 |
| LRRK2 | NMC | - | 111 | 0.52 | 0.63 | 0.55 | 0.37 | 115 | 0.30 | 0.31 | 0.29 | 0.10 |
|  | PD | - | 110 | 0.52 | 0.61 | 0.50 | 0.41 | 117 | 0.32 | 0.33 | 0.32 | 0.11 |
|  | - | G2019S | 203 | 0.53 | 0.64 | 0.54 | 0.40 | 216 | 0.30 | 0.32 | 0.30 | 0.11 |
|  |  | R1441G | 18 | 0.37 | 0.42 | 0.38 | 0.22 | 16 | 0.32 | 0.33 | 0.32 | 0.09 |
| GBA | NMC | - | 91 | 0.93 | 2.2 | 1.0 | 6.1 | 87 | 0.52 | 0.84 | 0.58 | 1.5 |
|  | PD | - | 60 | 0.75 | 1.6 | 0.87 | 2.4 | 91 | 0.51 | 1.0 | 0.60 | 2.1 |
|  | - | E326K | 7 | 0.78 | 0.79 | 0.60 | 0.55 | 9 | 0.49 | 0.45 | 0.42 | 0.17 |
|  |  | T369M | 5 | 0.73 | 0.62 | 0.56 | 0.27 | 6 | 0.37 | 0.37 | 0.37 | 0.07 |
|  |  | N370S | 119 | 0.82 | 1.1 | 0.84 | 1.0 | 111 | 0.49 | 0.59 | 0.51 | 0.67 |
|  |  | L444P | 8 | 0.92 | 1.0 | 0.73 | 0.82 | 6 | 0.54 | 0.50 | 0.49 | 0.12 |
|  |  | N370S/N370S | 12 | 6.1 | 12.2 | 7.3 | 14.3 | 11 | 5.1 | 6.2 | 4.4 | 4.6 |
| LRRK2/GBA | - | G2019S/N370S | - | - | - | - | - | 9 | 0.52 | 0.54 | 0.52 | 0.16 |

While PPMI has strong representation of G2019S LRRK2 and N370S GBA mutations, the cohorts for this study also included other more common risk and pathogenic variants of interest within these genes. Thus, we next assessed GlcSph levels at the variant level. Since disease status had no effect on either central or peripheral sources of GlcSph within LRRK2 and GBA carriers, we combined the NMC and PD group within each respective variant. CSF GlcSph was significantly increased in all GBA variants compared to non-carrier healthy controls (Fig. 1C). The degrees of elevation correlated with biochemical variant severity in some cases, with the smallest elevation observed in the heterozygous T369M common risk variant (93%, p=0.041) to the highest elevation observed in the heterozygous L444P pathogenic variant (163%, p=9.40e-05; Fig. 1C). However, no significant differences were observed between E326K, N370S, and L444P variants when comparing these groups to one another, suggesting that mutation severity does not necessarily correlate with CSF GlcSph level (Fig. 1C). Plasma GlcSph in the T369M variant was not statistically different from non-carrier healthy controls whereas the E326K variant had a 35% increase compared to non-carrier healthy controls that reached statistical significance (p=0.03; Fig. 1D). There were no statistical differences in plasma GlcSph when comparing between heterozygous E326K, N370S, and L444P variants (Fig. 1D). When comparing the allelic dosage of the N370S GBA mutation on CSF GlcSph levels, we found that the homozygous group was 841% higher than the heterozygous group (p=2.17e-31; Fig. 1C). This was consistent with plasma GlcSph, as there was an 813% increase in homozygous N370S carriers compared to heterozygous (p=1.62e-85; Fig. 1D). When focusing on the LRRK2 variants, CSF GlcSph in the G2019S carriers was significantly increased compared to non-carrier healthy controls (36%, p=0.0002; Fig. 1C). Interestingly, this was not observed in plasma GlcSph, suggesting the relationship between G2019S and GlcSph may be CNS-specific (Fig. 1D). There was no difference in the ROC-domain pathogenic variant R1441G compared to non-carrier healthy controls, although there was a 37% increase (though not statistically significant, p=0.066) when comparing G2019S to R1441G (Fig. 1C). In addition, our plasma cohort contained carriers of both G2019S LRRK2 and N370S GBA mutations. This group also had elevated GlcSph compared to non-carrier healthy controls, which was comparable to the heterozygous GBA mutation alone (70%, p=4.32e-5; Fig. 1D). The summary of all unadjusted absolute values of each variant can be found in Table 4.

Within the CSF and plasma cohorts, 460 subjects had matching CSF and plasma samples, allowing us to correlate CSF and plasma GlcSph levels within-subject. In GBA carriers, we observed a modest Pearson correlation of 0.54 within the GBA-PD group, whereas GBA-NMCs did not show any correlation between CSF and plasma GlcSph levels (Fig. 1E). No correlation was found between CSF and plasma GlcSph in the LRRK2 carriers nor non-carrier group regardless of disease status (Suppl. Fig. 1A-B).

### CSF GCase activity is decreased in sporadic PD and GBA carriers

We next analyzed GCase activity in the CSF and plasma using the 4-MU assay in 640 CSF samples (collected from 116 HC, 110 sPD, 120 LRRK2-NMC, 120 LRRK2-PD, 104 GBA-NMC, 70 GBA-PD) and 628 plasma samples (collected from 112 HC, 106 sPD, 115 LRRK2-NMC, 117 LRRK2-PD, 87 GBA-NMC, 91 GBA-PD). As expected, CSF and plasma GCase activity were significantly decreased in the GBA-NMC group by 33% and 13%, respectively, compared to healthy controls (p<0.0001 and p=0.0002), and by 30% and 13%, respectively when omitting the N370S homozygous carriers (p<0.0001 and p=0.0004;Fig. 2A, B). We also observed an 11% decrease in CSF GCase activity in the sPD group compared to healthy controls, while no difference was observed in plasma between these two groups (Fig. 2A, B). In addition, there was no statistical difference between LRRK2-NMCs and healthy controls in both biomatrices (Fig. 2A, B). There were no differences observed between NMC and PD subgroups within each respective monogenic cohort.

**Figure 2.**
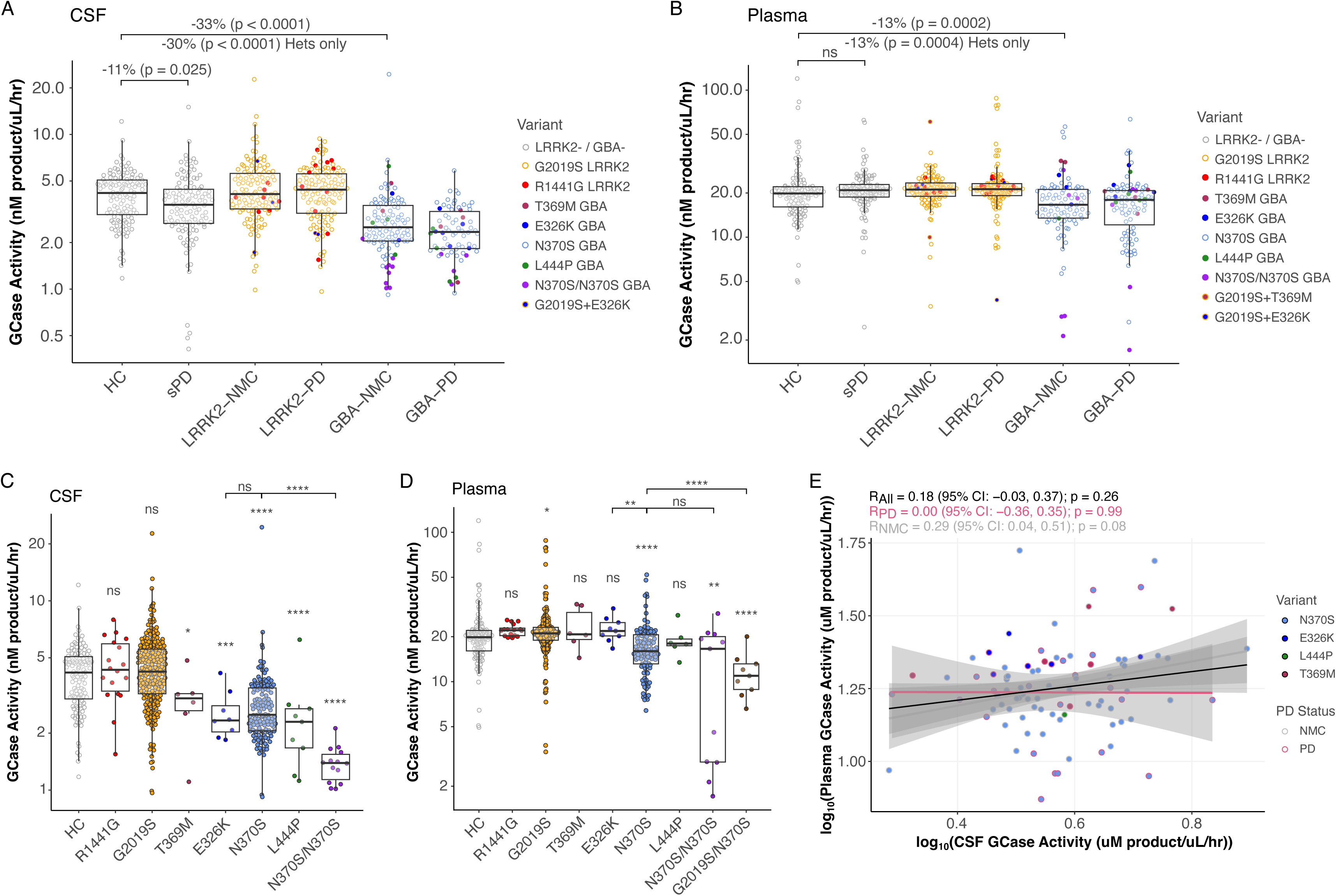
CSF and plasma GCase activity are reduced in GBA mutation carriers and sporadic Parkinson’s disease. **A** CSF and **B** plasma glucocerebrosidase (GCase) activity measured by the 4-methylumbelliferyl-β-D-glucoside (4-MU) assay across healthy controls (HC), sporadic Parkinson’s disease (sPD), LRRK2 non-manifesting carriers (LRRK2-NMC), LRRK2-PD, GBA non-manifesting carriers (GBA-NMC), and GBA-PD. **C** CSF and **D** plasma GCase activity stratified by individual genetic variants. The asterisks above each boxplot indicate statistical significance relative to HCs unless otherwise specified by brackets. **E** Correlation between paired CSF and plasma GCase activity within GBA carriers. Data are shown as boxplots with individual data points overlaid; boxes represent the interquartile range with median indicated, and whiskers denote 1.5× IQR. Statistical comparisons were performed using ANCOVA pairwise tests with Benjamini–Hochberg correction for multiple comparisons; adjusted *p*-values are shown. Correlations were assessed using Pearson correlation on log-transformed values. Asterisks denote significance; * p < 0.05, ** p < 0.01, *** p < 0.001.

When stratified by variant type, GCase activity was decreased in accordance with GBA mutation severity in both CSF and plasma (Fig. 2C, D). Interestingly, when focusing on the common PD risk variants T369M and E326K, there were significant decreases in CSF GCase activity compared to healthy controls (-34%, p=0.01 and –40%, p=0.0003, respectively) whereas both of these variants were comparable to healthy controls in the plasma (Fig. 2C, D). In the CSF, no difference was observed between E326K and heterozygous N370S, whereas a significant difference was observed between these variants in the plasma (25%, p=0.004). CSF GCase activity decreased 48% in the homozygous N370S carriers compared to heterozygous carriers of the same variant, suggesting a linear relationship between allelic dosage and GCase loss of function (p=7.14e-10; Fig. 2C). Interestingly, this was not the case for plasma GCase activity, where no difference was found between these groups (Fig. 2D). When analyzing LRRK2 variants, CSF GCase activity was similar to healthy controls in both the G2019S and R1441G groups, while plasma GCase activity was significantly increased in the G2019S carriers (8.6%, p=0.01; Fig. 2C) which was consistent with GCase activity measured in dried blood spots in a previous study^19^. In addition, there was a 42% reduction in GCase activity in the individuals harboring both a G2019S LRRK2 and a N370S GBA mutation compared to healthy controls (p=6.47e-9; Fig. 2D). Surprisingly, this was significantly lower than the heterozygous N370S group alone by 32% (p=7.63e-5; Fig. 2D). Finally, when correlating GCase activity within GBA carriers with matching CSF and plasma samples collected from the same visit, there was no correlation found regardless of disease status (Fig. 2E). Similarly, no correlations were observed in either the LRRK2 carriers or in the non-carrier subgroup (Suppl. Fig. 1C-D). Additionally, we correlated GlcSph level and GCase activity within each genetic subgroup of each biomatrix. Regardless of biomatrix, disease status, or genetic status, there were no correlations observed between GlcSph levels and GCase activity (Suppl. Fig. 2).

### CSF and plasma GlcSph/GCase activity are differentially affected by age and sex

We next looked for differences and correlations between GlcSph levels or GCase activity with demographics like sex and age, respectively. Using ANCOVA pairwise tests between males and females per genetic and nongenetic subgroup, we observed a statistically significant difference between sex and CSF GlcSph in the LRRK2-PD, non-carrier healthy control and sporadic PD groups after adjusting for age in which males had higher levels of CSF GlcSph than females (unadjusted p=0.003, 0.02, and 0.02, respectively; Fig. 3A). We also observed that males had lower levels of CSF GCase activity than females in the GBA-PD subgroup (p=0.01; Fig. 3B). Overall, regardless of subgroup we observed a similar pattern across all CSF measurements in which males had higher levels of GlcSph and lower levels of GCase activity compared to females, even though most groups did not reach statistical significance (Fig. 3A-B). Consistent with prior findings of reduced ceramide levels in males with sPD^20^, these results suggest increased GCase pathway burden in males, potentially contributing to the higher prevalence of PD in men (∼1.5x;^21,22^). Additionally, there were no differences observed in any plasma measurements (Suppl. File 1). Next, partial Pearson correlations adjusted for sex were used to determine relationships between these measurements and age. We observed an overall positive correlation between CSF GlcSph and age (Fig. 3C-E), where most subgroups showed modest but significant positive correlations with age ranging from R=0.36 to 0.27 (Fig. 3C-E; Suppl. File 1). In addition, a weak negative correlation was observed between plasma GCase activity and age (Fig. 3F-G). Interestingly, when stratified by disease status, this finding was largely driven by those without PD, with the non-carrier healthy control group having a R = -0.19 and the LRRK2-NMC group having a R = -0.24 (Fig. 3F-G) of which statistical significance was lost after Benjamini-Hochberg correction (Suppl. File 1). The GBA carrier subgroup did not show any significant correlation between plasma GCase activity and age (Fig 3H; Suppl. File 1). There were also no significant correlations between CSF GCase activity nor plasma GlcSph and age (Suppl. File 1).

**Figure 3.**
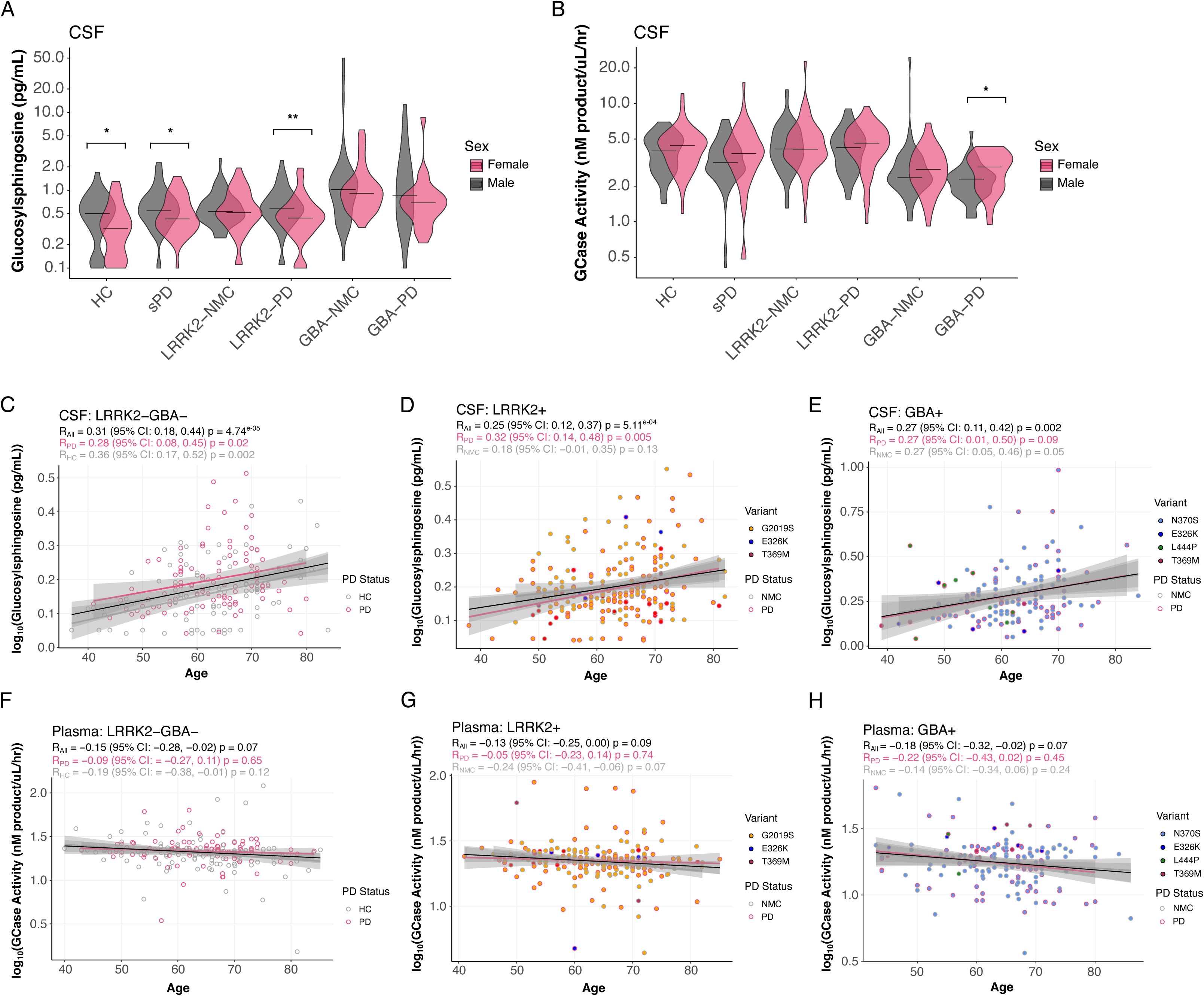
CSF GlcSph and plasma GCase activity show sex- and age-dependent associations across Parkinson’s disease subgroups. Violin plots depict sex-stratified distributions of **A** CSF GlcSph and **B** CSF GCase activity across diagnostic groups with unadjusted p-value significance displayed. Partial Pearson correlations adjusted for sex are shown between age and CSF GlcSph for **C** non-carriers, **D** LRRK2 carriers, and **E** GBA carriers. Correlations between age and plasma GCase activity are shown for **F** non-carriers, **G** LRRK2 carriers, and **H** GBA carriers. Benjamini–Hochberg corrections were applied for multiple comparisons; significance is denoted within each panel. Homozygous N370S mutation carriers were omitted from the statistical analyses to prevent correlations driven by this group. Asterisks denote significance; * p < 0.05, ** p < 0.01, *** p < 0.001.

### GlcSph and GCase activity measurements show association with clinical assessments and other PD GWAS variants

We also generated partial Spearman correlations in participants with PD across a selection of clinical assessments available through PPMI. Since individuals with GBA-PD are known to have a more aggressive form of the disease based on an earlier age at onset and worse cognitive impairments compared to those with non-GBA-PD^23,24^, we included several motor, cognitive, and psychological outcomes. After adjusting for sex, age, and PD medication status, the most robust correlation found was between plasma GlcSph and UPSIT score within the GBA-PD group, demonstrating that higher plasma GlcSph correlates with worse UPSIT scores (rho = 0.48; unadjusted p=0.0007) (Fig. 4A, C). Similar trends were also observed with CSF GlcSph in the GBA-PD group, although not statistically significant (rho = 0.21; unadjusted p=0.12; Fig. 4A-B). LRRK2-PD CSF and plasma GlcSph groups had no correlation with UPSIT score regardless of biofluid (Fig. 4A-C). This remained true when stratifying the group on SAA status (data not shown). While the sporadic PD group did not demonstrate a significant correlation between UPSIT score and GlcSph levels in either biofluid (Fig. 4A), boxplot visualization indicated a graded increase in CSF GlcSph with increasing severity of olfactory impairment, similar to the trend observed in the GBA-PD group (Fig. 4B). Of all other assessments, MDS-UPDRS Part 3 total score and its subgroupings based on symptom had the most correlations with CSF and plasma GlcSph across the genetic subgroups, although none particularly showed significant correlation across the genetic subgroups consistently (Fig. 4A). Overall, there were more hits of statistical significance when correlating clinical assessments with GlcSph compared to GCase activity. The heatmap shown in Figure 4A displays all clinical assessments where at least one subgroup showed significant correlation with GlcSph and/or GCase activity. The full results of all clinical assessments tested can be found in Supplementary File 2.

**Figure 4.**
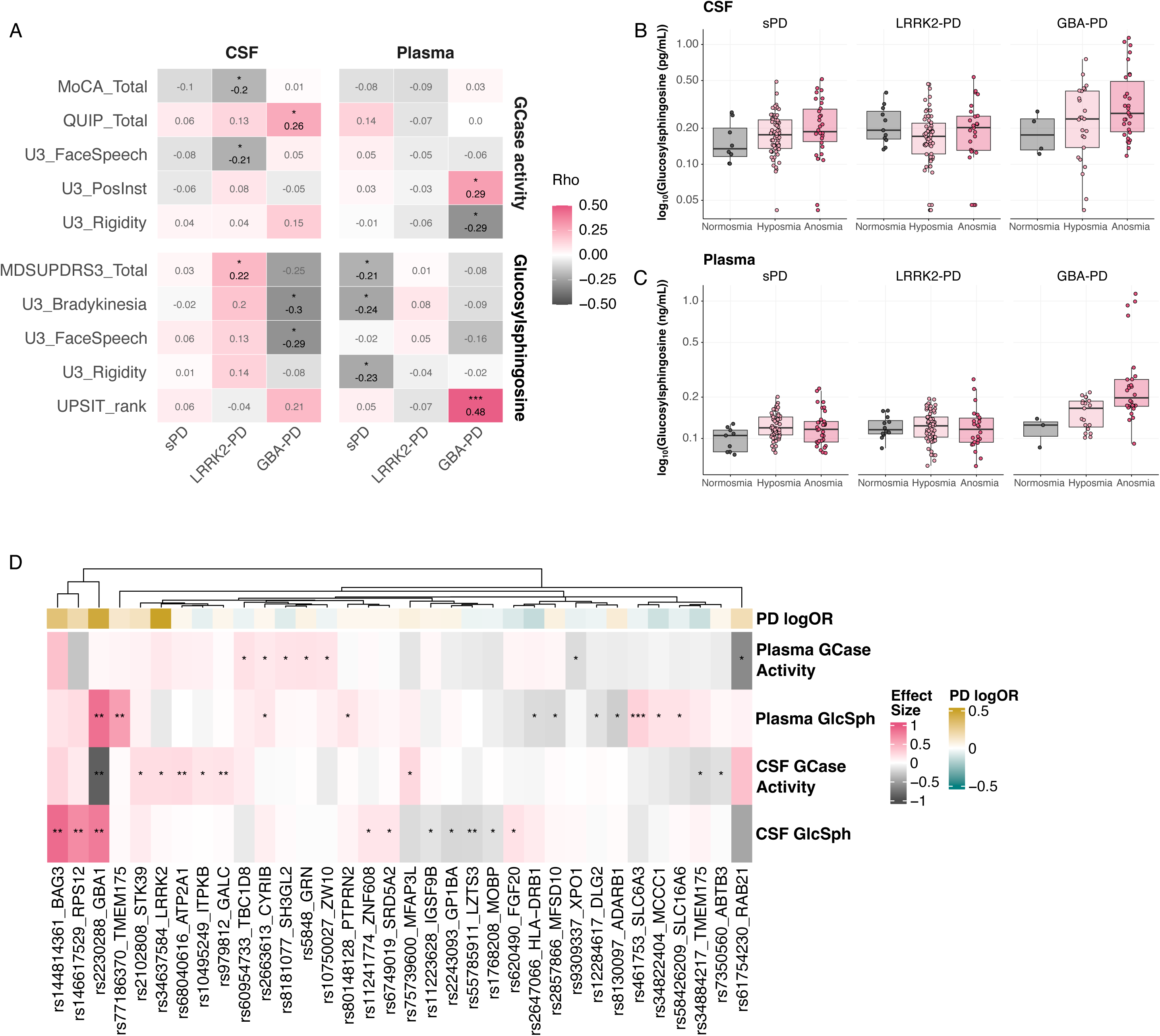
Associations of GlcSph and GCase activity with clinical assessments and PD-associated variants. **A** Partial Spearman correlation heatmap summarizes associations between CSF and plasma GlcSph levels and GCase activity and clinical assessments in participants with PD and stratified by genetic subgroup. Correlations were adjusted for age, sex, and PD medication status. Color scale represents Spearman’s rho, and statistical significance is indicated within each cell using asterisks (unadjusted p-values). Correlation coefficients are also annotated within each cell, with grey text denoting ‘not significant’ correlations. MoCA: Montreal Cognitive Assessment; QUIP: Questionnaire for Impulsive-Compulsive Disorders in Parkinson’s Disease; U3_FaceSpeech: MDS-UPDRS Part 3 hypomimia and hypokinetic dysarthria score; U3_PostInst: MDS-UPDRS Part 3 postural instability score; U3_Rigidity: MDS-UPDRS Part 3 rigidity score; MDSUPDRS3_Total: Movement Disorder Society-Unified Parkinson’s Disease Rating Scale Part 3 total score; U3_Bradykinesia: MDS-UPDRS bradykinesia score; UPSIT_rank: University of Pennsylvania Smell Identification Test. **B** CSF and **C** Plasma GlcSph levels stratified by olfactory status (normosmia, hyposmia, anosmia). Data are shown as boxplots with individual data points overlaid; boxes represent the interquartile range with median indicated, and whiskers denote 1.5× IQR. **D** Heatmap of associations between PD-associated variants and CSF and plasma GCase activity and GlcSph levels. Variants depicted are those from the 140 analyzed that resulted in at least one significant (p < 0.05) association. Statistics generated from linear models of log-transformed analytes adjusted for sex, age, and genetic ancestry principal components and subject to inverse-normal transformation. Effect sizes are in units of SD change of adjusted and transformed analyte levels per dosage of PD-associated allele, either risk or protective as indicated by the log odds ratio in the column annotation. Asterisks denote significance; * p < 0.05, ** p < 0.01, *** p < 0.001.

We next examined the genetic determinants of these analytes, restricting analyses to established PD-associated variants given the limited sample size to identify SNP–biomarker associations. Carriers of pathogenic *GBA* mutations were excluded, as these variants are expected to exert strong effects on GlcSph and GCase activity and could disproportionately influence the results. As a positive control, common *GBA* risk variants were retained. Overall, 34 PD-linked variants demonstrated association with at least one of CSF or plasma GlcSph or GCase activity levels. The strongest associations were observed at the *GBA* locus, with the PD risk allele at the E326K variant (rs2230288) showing association with elevated CSF and plasma GlcSph levels and reduced CSF GCase activity (p = 0.0064, p = 0.0015, and p = 0.0015, respectively; Fig. 4D; Suppl. File 3). Beyond *GBA*, the next strongest associations were detected between CSF GlcSph levels and variants at the *BAG3* and *RPS12* loci (p = 0.0037 and p = 0.0070, respectively; Fig. 4D; Suppl. File 3). Notably, variants exhibiting the strongest associations with GlcSph and GCase activity also correspond to loci with relatively large PD (log) odds ratios, suggesting that dysregulation in these lysosomal biomarkers is not limited to rare mutations in *GBA* itself but rather extends to more common PD genetic risk factors across the genome.

## Discussion

Reliable biomarkers are critical in Parkinson’s disease for early diagnosis, monitoring disease progression, and assessing pharmacodynamic responses that can inform the development of targeted therapies. This study provides the first large-scale demonstration that CSF GlcSph is elevated not only in GBA mutation carriers but also in sporadic PD and LRRK2 G2019S carriers, indicating shared central GCase pathway perturbation across PD subtypes. Within the GBA carrier groups, elevated GlcSph levels were observed not only in carriers of pathogenic GBA mutations but also in individuals harboring the common GBA risk variants E326K and T369M. These findings indicate that PD-specific GBA risk variants are associated with central GCase pathway dysfunction despite not being traditionally classified as pathogenic mutations. This observation suggests that biological evidence of pathway perturbation, rather than mutation classification alone, may be more informative for identifying patients likely to benefit from GCase-targeted therapies. However, replication in larger cohorts of common variant carriers will be necessary to confirm this finding. Similarly, among LRRK2 carriers, elevated GlcSph was observed primarily in G2019S carriers, whereas R1441G carriers did not differ significantly from non-carrier healthy controls. Although the limited number of R1441G carriers restricts interpretation, this finding raises the possibility that kinase-domain and GTPase-domain LRRK2 variants differentially influence lysosomal biology and GCase pathway function. Together, these results highlight the importance of considering variant-specific biological effects when evaluating lysosomal biomarkers and selecting patient populations for precision medicine approaches in PD.

Notably, our data further demonstrate that the presence of a *GBA* mutation alone does not predict GlcSph accumulation. Several individuals harboring pathogenic *GBA* variants exhibited GlcSph levels comparable to healthy controls, indicating that reduced GCase activity is not sufficient, in isolation, to drive substrate accumulation. This disconnect suggests that additional modifiers—such as lysosomal capacity, lipid metabolism, or cellular stress responses—likely determine whether GCase deficiency translates into measurable glycosphingolipid buildup. In this context, GlcSph may represent a more proximal indicator of pathogenic burden than GCase activity alone. The requirement for additional biological factors to drive GlcSph accumulation may help explain the incomplete penetrance of *GBA* mutations in Parkinson’s disease, whereby only a subset of mutation carriers develop clinically manifest disease.

Moreover, poor correlation between central and peripheral GlcSph levels across groups was observed, with only a significant positive correlation detected in the GBA-PD subgroup. This lack of concordance likely reflects fundamental biological differences between compartments that may lead to GlcSph accumulation through compartment-specific mechanisms of action and thus may be the reason that *GBA* mutations can cause peripheral and central diseases, namely GD and PD, respectively. This finding suggests that CSF is the more relevant biofluid when evaluating GlcSph burden in PD. Dopaminergic neurons are uniquely vulnerable to oxidative stress and therefore highly dependent on intact lysosomal function^25^. Perturbation of the GCase pathway within the CNS may preferentially affect these neurons by exacerbating lysosomal dysfunction, glycosphingolipid accumulation, and impaired α-synuclein clearance. These findings further indicate that plasma GlcSph is unlikely to serve as a reliable surrogate for CNS GlcSph burden. Clinically, this implies that lumbar punctures would be necessary to understand the degree of GCase-pathway perturbation in the CNS of patients with PD and when evaluating pharmacodynamic response in therapeutics targeting the CNS.

GCase activity was reduced in *GBA* mutation carriers in both biomatrices, with the magnitude of reduction reflecting variant severity and allelic dosage, consistent with prior biochemical and genetic studies^19,26,27^. We also observed a modest but significant reduction in CSF GCase activity in sporadic PD, supporting the notion that GCase pathway dysfunction extends beyond genetically defined GBA-associated PD. In contrast, plasma GCase activity showed distinct behavior, including age-dependent decline in non-carrier and LRRK2 non-manifesting carriers and increased activity in G2019S LRRK2 carriers that was not observed in the CSF, underscoring the biochemical independence of central and peripheral compartments. Importantly, despite robust group-level differences, GCase activity did not correlate with GlcSph levels within either CSF or plasma. This lack of correlation likely reflects fundamental differences in biological timescale and compartmental regulation, as the enzyme activity assay captures instantaneous catalytic capacity under optimized *in vitro* conditions of extracellular GCase, whereas GlcSph concentrations represent a steady-state integration of substrate flux, cellular turnover, and compensatory mechanisms over time. Thus, these measures of GCase activity and GlcSph levels may provide complementary but non-redundant information about GCase pathway dysfunction, and the absence of direct correlation does not diminish the value of either measure as a biomarker.

Currently, the PD field lacks well-established biomarkers of disease and progression, making it challenging to predict clinical severity and to stratify patients for disease-modifying therapies. Among the clinical measures examined, the strongest association was observed between GlcSph and hyposmia, a feature that frequently precedes motor symptom onset in PD. Given the exploratory nature of this analysis, replication of this association in another cohort will be necessary to verify this finding as well as inclusion of longitudinal data, as the current study only evaluates associations from cross-sectional samples.

While the number of disease-modifying therapeutics targeting genetic mutations in PD has increased in recent years, making up 44% of the total active clinical trials for PD in 2024^28^, improving the chance of clinical success requires the thoughtful selection of an appropriate patient population, as well as selection of the appropriate dose, treatment regimen, and pharmacodynamic biomarkers. Patient selection in the context of PD requires addressing and tailoring therapies to genetic and biological phenotypes within the broad heterogeneity often observed in PD. In this framework, CSF GlcSph may also enable identification of GBA-like phenotypes within the sporadic PD population, extending the applicability of GCase-targeted therapies beyond genetically defined subgroups. Evaluating CSF GlcSph in ongoing and future trials targeting the GCase pathway will be essential to validate its utility as a target engagement biomarker and to define its relationship to dose, treatment duration, and clinical response. Integration of CSF GlcSph with complementary biomarkers of lysosomal function, α-synuclein pathology, and neurodegeneration may further clarify how GCase pathway perturbation interfaces with broader disease mechanisms. Additionally, expanding analyses to include diverse ancestries, additional lysosomal gene variants, and independent cohorts beyond PPMI will be important to assess generalizability and variant-specific effects. Together, these efforts will help define the role of CSF GlcSph as a biomarker for Parkinson’s disease and guide rationale for patient stratification and dose selection in future disease-modifying trials.

## Methods

### Cohort

Samples were obtained through PPMI to investigate the lipidome and metabolome of individuals with monogenic and sporadic PD (Project ID #180). After the completion of this study, leftover volumes were used to generate the current study data to assess GlcSph and GCase activity in CSF and plasma (Project ID #303). Samples were chosen based on age and sex balancing. Samples collected at the baseline visit for both CSF and plasma were prioritized in addition to annual follow up collections at 1 and 2 years for plasma samples. A detailed report of the demographics and other sample characteristics are outlined in Table 1 and Table 2, respectively. All data are publicly available through PPMI’s online database (https://ida.loni.usc.edu). For more information on the PPMI study, detailed protocols for participant selection, clinical assessment, and data collection have been described previously^29^.

### Glucosylsphingosine assay

#### Assay qualification specifications

The targeted GlcSph assays for CSF and plasma are qualified assays with a dynamic range of 0.1 - 50 pg/mL and 0.1 - 10 ng/mL, respectively, using an AB Sciex 7500 or an AB Sciex 7500+ mass spectrometer. The assay qualification has undergone precision and accuracy runs, benchtop stability, and freeze-thaw stability tests in each matrix. Long-term stability is currently being assessed in both matrices, with up to 504 days for CSF and 721 days for plasma passed at the time of manuscript preparation. Qualification specifications are summarized in Table 3.

Since the average age of CSF samples from our PPMI cohort is around 10 years old, we first ran a small validation study to determine whether we could detect CSF GlcSph in this cohort by running 6 technical replicates of a pooled CSF sample. This pooled sample was created by pooling 10 µL from each of the 640 total CSF samples in our parent cohort. We also performed a dilution linearity test of these samples at 2-, 5-, 10-fold dilutions. Samples were within the detectable range at all three dilution factors. CVs at each factor were 1.4%, 2.6%, and 1.7% with an accuracy of 102.3%, 99.8%, and 95.6%, respectively. The concentration of pooled CSF from PPMI samples was 1.8 pg/mL, ensuring that the average GlcSph levels in our PPMI samples were within the detectable range of the assay.

### Project ID #303

#### CSF Sample preparation and protocol

For CSF samples, the GlcSph LC-MS/MS assay was performed in ten equally distributed batches based on repository site, gender, disease status, and genetic status. These batches were divided into two rounds (4 batches in round 1 and 6 batches in round 2). This assay requires a volume of at least 100 µL per sample, thus, not all samples from the original parent cohort had sufficient volume leftover for GlcSph analysis. As a result, there are only n=568 samples analyzed from the original n=640. A total of 2 pooled control and PD CSF sample replicates were created and distributed evenly within and between batches to enable evaluation of inter-batch deviations. Values below the quantification limit (BQL) were imputed by half of the lower limit of quantification (LLOQ) reported. To incorporate as many samples as possible, the LLOQ varied based on the dilution of each CSF sample, thus, BQLs <0.2 with 2-fold dilution (50 µL sample), <0.222 with 2.22-fold dilution (45 µL sample), and <0.25 with 2.5-fold dilution (40 µL sample) were imputed as 0.1, 0.111, and 0.125, respectively. All samples were run in a blinded.

To prepare samples for LC-MS/MS analysis, aliquots of 100 μL of calibration standards (STDs), quality control samples (QCs) in methanol, and experimental CSF samples were transferred to a clean 2-mL protein LoBind 96-well plate. 100 μL of artificial CSF (aCSF) (Tocris Bioscience cat # 3525) was added to the STDs and QCs while 100 μL of methanol was added to the CSF samples and CSF matrix QCs. 20 μL of internal standard working solution (10 pg/mL of GlcSph-d5 (Avanti Polar Lipids cat # 860636P-1mg) in methanol) was added to each sample except blank samples while 20 μL of methanol was added to the blank samples. 800 μL of methanol/water/formic acid (450:550:5, v/v/v) was added to all samples. After mixing, the samples were loaded onto a 2.0 mL Strata-X Polymeric Reversed Phase plate (Phenomenex cat # 8E-S100-TGB), which was preconditioned with 1mL of hexane twice followed by 1 mL of methanol twice and equilibrated with 1 mL of methanol/water/formic acid (450:550:5, v/v/v) twice. Once samples were completely absorbed in the plate sorbent, the plate was washed with 1 mL of methanol/water/formic acid (450:550:5, v/v/v) twice and 1 mL of methanol/water (450:550, v/v). After the washing, 1 mL protein LoBind 96-well collection plate was placed under the extraction plate. The analyte was eluted by 400 μL of methanol/ammonium hydroxide (25%) (100:0.04, v/v) twice into a collection plate. The eluent was dried down completely under purified nitrogen gas flow at 60 °C and reconstituted in 120 μL acetonitrile/methanol/water/formic acid/ammonium formate (1M) (2850:75:60:3:15, v/v/v/v/v) before the samples were injected to LC-MS/MS for analysis.

LC-MS/MS analysis were performed on an Exion LC AD UHPLC system coupled with a Sciex API 7500 or 7500+ Triple Quad mass spectrometer (AB Sciex, Framingham, MA). The MRM transitions for the analyte and internal standard were 462.3 to 282.1 and 467.6 to 287.4, respectively. Collisional Energy was 27 eV. HPLC chromatography was established on a Halo, 90 Å, HILIC column (2 μm, 3.0 x 150 mm) (Advance Materials Technology cat # 91813-701) and the column was kept at 35 °C during the run. For the LC separation of GlcSph from the matrix interference, the two mobile phases used were 0.1 % Formic Acid and 50 mM ammonium formate in water (mobile phase A) and 0.1 % Formic Acid in acetonitrile/methanol (95:5, v/v) (mobile phase B). The flow rates were 0.5 mL/min for analyte elution and 0.3 mL/min for column wash and equilibration. Mobile phase B concentration was initially set at 91% and kept for 8.5 min, and then decreased to 20% at 8.6 min and kept until 10.60 min before increasing mobile phase B concentration to 91% at 10.70 min.

#### Plasma sample preparation and protocol

For plasma samples, the GlcSph LC-MS/MS assay was performed in twenty-two equally distributed plates based on repository site, gender, disease status, and genetic status. These batches were divided into two rounds (18 batches in round 1 and 4 batches in round 2). This assay requires a volume of 20 µL per sample and performed in the in all plasma samples (n=640). A total of 2 matrix QC sample replicates were created and distributed evenly within and between batches to enable evaluation of inter-batch deviations. For these runs, the BLQ was <0.1 ng/mL and the ALQ was >10 ng/mL. All samples were run blinded.

To prepare samples for LC-MS/MS analysis, aliquots of 20 μL of STDs, QCs in methanol, and experimental plasma samples were transferred to a clean glass 1-mL 96-well plate. 20 μL of 1% bovine serum albumin (BSA) was added to the STDs and QCs while 20 μL of methanol was added to the plasma samples and plasma matrix QCs. 10 μL of internal standard working solution (5 ng/mL of GlcSph-d5 in Halo B (0.5% Formic and 5 mM Ammonium Formate in Acetonitrile/Isopropanol/H_2_O (92.5/5/2.5))) was added to each sample except blank samples while 10 μL of Halo B was added to the blank samples. 150 μL of 1% ammonium hydroxide in water was added to all samples. After mixing, the samples were loaded onto a 200 μL Isolute SLE+ (Biotage cat # 820-0200-P01), which was placed on a clean glass 1-mL 96-well plate. ∼6 psi positive pressure was applied onto the SLE plate for approximately 30 seconds or until all the samples are loaded onto the plate. The analyte was eluted by adding 1000 μL of ethyl acetate. The eluent was dried down completely under purified nitrogen gas flow at 40 °C and reconstituted in 150 μL Halo B before the samples were injected to LC-MS/MS for analysis. LC-MS/MS conditions for plasma samples were the same as those for CSF samples.

#### GCase assay – sample preparation and protocol

To ensure that independent factors and covariates of interest were evenly distributed among batches and randomized, both CSF and plasma samples were equally distributed into batches based on repository site, gender, disease status, and genetic status. All samples were run blinded.

To measure the endogenous activity of GCase in plasma and CSF, we employed the well-established 4-methylumbelliferyl β-D-glucopyranoside (4-MUG) assay (Sigma Aldrich, Cat. #M3633). In this assay, synthetic substrate, 4-MUG, is added to each sample to determine rate of hydrolysis by GCase into the fluorescent byproduct 4-MU.

For CSF samples, a 400-fold dilution of 500mM 4-MUG substrate into GCase activity buffer was performed. 100µL of the mixed buffer containing substrate was then aliquoted into the wells of a 96-well V-bottom plate and 25uL of CSF was added to each well. For plasma samples, a 30-fold dilution of 30mM 4-MUG substrate into GCase activity buffer was performed. After mixing, 129uL was dispensed into each well of a 96-well V-bottom plate and 4uL of plasma was added. Six pooled control samples were also added to each plate in order to calculate inter-assay reliability as well as negative control samples treated with Conduritol B Epoxide (CBE). The final concentration of 4-MUG was 1mM. Samples were then pipetted to mix and 100uL were transferred into a Corning® 96-well black round bottom polypropylene plate. Plates were sealed and left to shake at room temperature for 10 minutes followed by incubation overnight at 37°C without carbon dioxide. On the following day, plates were spun at 1000 g for 60 seconds to precipitate condensation formed on the top of the seal and sides of the wells. Standard curve samples of 4-MU in GCase activity buffer were added to each plate. Then 100uL per well of GCase stop buffer was added. Plates were read on a SpectroMAX using Ex 365/Em 445.

To transform raw data into nM product/uL/hr, a linear model was fit after adjusting for batch number. Mean residuals derived from this linear model were used to correct for batch-dependent variances in the log transformed sample measurement data. Following batch correction, data were back transformed to linear scale and a 4-MU standard curve was used to interpolate the arbitrary unit values into the concentration of 4-MU produced per sample. To get a unit of substrate product produced over time, we divided the interpolated value by the sample volume and incubation time. Using this methodology, we derived the final unit of “nM product/uL/hr”.

#### Statistical analyses

To analyze genetic group-level and variant-level differences between healthy controls, sPD, and monogenic groups, we fit ANCOVA robust linear regression models adjusted for age, sex, batch effect, and standard of care medication, namely levodopa and MAOB inhibitors, of log2 transformed values. For the measurements CSF and plasma GlcSph, values that were below the level of quantification were set to one half of the LLOQ of each assay. Values that were higher than the upper limit of detection were multiplied by 1.25x the ULOQ of each assay. For the group-level pairwise comparisons, we omitted the homozygous N370S carriers to prevent overestimation of statistical differences. Pearson correlations were calculated within subject between CSF and plasma of each analyte within each genetic group. Pearson correlations were also calculated within-subject to assess the relationship between GlcSph levels and GCase activity of each biomatrix and genetic group. Homozygous N370S carriers were omitted from these analyses as they drastically affected correlation outcome in the GBA+ groups. To analyze sex differences within each genetic and nongenetic subgroup, ANCOVA robust linear regression models adjusted for age of log2 transformed values were used. For all partial Pearson correlations, age, sex and PD standard of care medications were treated as covariates where specified. A Fisher z *k*=1 covariate was used to calculate 95% confidence intervals for the partial Pearson correlations for age. Both Pearson, partial Pearson, and partial Spearman correlations were calculated on log10-transformed values. All p-values reported are Benjamini-Hochberg adjusted unless otherwise stated. To assess correlations with MDS-UPDRS Part 3 scores, scores recorded during the OFF state were used when specified. When multiple scores were recorded per visit without ON/OFF state specification, the highest, worst performing score was counted. If a single score was recorded within a given visit regardless of ON/OFF state specification, this score was used in the analysis. In addition to assessing total score of part 3, five subgroups were made and assessed individually based on symptom domains: facial expression/speech (questions 3.1-3.2), rigidity (3.3), bradykinesia (3.4-3.8, 3.14), postural instability (3.9-3.13, and tremor (3.15-3.18). To assess correlation with UPSIT categorical rank, normosmia, hyposmia, and anosmia were coded as 1, 2, and 3, respectively.

#### Genetic association analyses

To understand the genetic contributions (beyond *GBA* and *LRRK2*) to levels of plasma and CSF GCase activity and GlcSph, we investigated the relationship between known PD-associated variants and levels of our analytes of interest in the profiled PPMI cohort. To define PD-associated variants, we used the list of 157 SNPs (representing 157 independent signals) across 134 loci as identified in the most recent PD GWAS study of European ancestry samples^4^. While genetic contributions to GCase activity and GlcSph levels may exist outside of these variants/regions, we sought to limit our investigation to this list of variants owing to our limited cohort size to discover SNP-to-analyte associations.

We utilized publicly available whole genome sequencing data on the profiled PPMI samples, as available through the Accelerating Medicines Partnership – Parkinson’s Disease (AMP-PD) data resource^30^ (2021_v2-5 release; N = 10,418 total and N = 1,807 PPMI subjects). As previously described^31,32^, we merged these data with reference samples from the 1000 Genomes Phase 3 and used the latter to predict genetic ancestry in the PPMI cohort. Subsequent analyses were restricted to those of European ancestry (N = 1,741) to match the ancestry of the PD GWAS data. We also removed GBA pathogenic variant carriers (e.g. N370S, L444P, etc.) in order to examine the effect of PD variants outside of these strong drivers of GCase activity and GlcSph. Finally, plasma measurements were restricted to those from the first available visit timepoint for that individual when repeated measures for an individual were present. After applying these filters, the following sample counts with both WGS data and analyte measurements were available: 422 for plasma GCase activity, 424 for plasma GlcSph, 442 for CSF GCase activity, and 396 for CSF GlcSph. Of the 157 lead PD GWAS SNPs, 140 were available in the PPMI WGS data.

As previously described^31,32^, to compute SNP to analyte statistical associations, we first regressed the natural log-transformed analyte measurements onto a linear model consisting of sex, age, and the first 3 principal components derived from the genome-wide WGS data from samples in the respective cohort. The residuals from this model were then inverse normal transformed using the blom() function from the “rcompanion” R package. The resulting values were used as outcomes variables in association testing against the 140 PD-associated SNPs using an additive model in plink 1.9^33^. We ensured that the tested allele in SNP-analyte statistics was identical to the tested allele in the PD GWAS, in order to interpret directionality of SNP to GCase activity and/or GlcSph changes with respect to the effect of the same allele on PD.

## Author Contributions

J.H.K. analyzed, visualized, and interpreted the data and conceptualized and drafted the manuscript. D.J. and R.R. ran the LC-MS/MS and 4-MU assays, respectively, and contributed to the methods section. B.T. and C.T. ran the LC-MS/MS assays. S.V.A. analyzed the relationship between analyte levels and PD GWAS SNPs and contributed to the respective methods and results sections. R.M. and E.A.M. participated in sample selection and study design and gave guidance for some of the statistical analyses reported in this manuscript. D.J., B.T., C.T., and M.F. developed and qualified the LC-MS/MS assays. A.G.H., M.A.S., and S.H-R. provided support for this study through sample acquisition, study design, and interpretation of findings. All authors read, edited, and approved the final manuscript.

## Acknowledgements

We want to thank the Parkinson’s Progression Markers Initiative for granting us the CSF and plasma samples analyzed in this study. We also thank the Michael J. Fox Foundation for the funding support of the shared grant between Denali Therapeutics and University of Harvard Medical School, MJFF-010325.

Data used in the preparation of this article were obtained from the Accelerating Medicine Partnership® (AMP®) Parkinson’s Disease (AMP PD) Knowledge Platform. For up-to-date information on the study, visit https://www.amp-pd.org. The AMP® PD program is a public-private partnership managed by the Foundation for the National Institutes of Health and funded by the National Institute of Neurological Disorders and Stroke (NINDS) in partnership with the Aligning Science Across Parkinson’s (ASAP) initiative; Celgene Corporation, a subsidiary of Bristol-Myers Squibb Company; GlaxoSmithKline plc (GSK); The Michael J. Fox Foundation for Parkinson’s Research; Pfizer Inc.; Sanofi US Services Inc.; and Verily Life Sciences. ACCELERATING MEDICINES PARTNERSHIP and AMP are registered service marks of the U.S. Department of Health and Human Services.

Data used in the preparation of this article was obtained on [2023-10-09] from the Parkinson’s Progression Markers Initiative (PPMI) database (www.ppmi-info.org/access-dataspecimens/download-data), RRID:SCR_006431. For up-to-date information on the study, visit www.ppmi-info.org. PPMI – a public-private partnership – is funded by the Michael J. Fox Foundation for Parkinson’s Research and funding partners, including 4D Pharma, Abbvie, AcureX, Allergan, Amathus Therapeutics, Aligning Science Across Parkinson’s, AskBio, Avid Radiopharmaceuticals, BIAL, BioArctic, Biogen, Biohaven, BioLegend, BlueRock Therapeutics, Bristol-Myers Squibb, Calico Labs, Capsida Biotherapeutics, Celgene, Cerevel Therapeutics, Coave Therapeutics, DaCapo Brainscience, Denali, Edmond J. Safra Foundation, Eli Lilly, Gain Therapeutics, GE HealthCare, Genentech, GSK, Golub Capital, Handl Therapeutics, Insitro, Jazz Pharmaceuticals, Johnson & Johnson Innovative Medicine, Lundbeck, Merck, Meso Scale Discovery, Mission Therapeutics, Neurocrine Biosciences, Neuron23, Neuropore, Pfizer, Piramal, Prevail Therapeutics, Roche, Sanofi, Servier, Sun Pharma Advanced Research Company, Takeda, Teva, UCB, Vanqua Bio, Verily, Voyager v. 25MAR2024 Therapeutics, the Weston Family Foundation and Yumanity Therapeutics.

## Disclosures

J.H.K, D.J., S.V.A., R.M., B.T., C.T., M.F., A.G.H., and S.H-R. are employees of and hold stock in Denali Therapeutics Inc. R.R. may hold stock in Denali Therapeutics Inc.

---

**Supplementary Figure 1.**
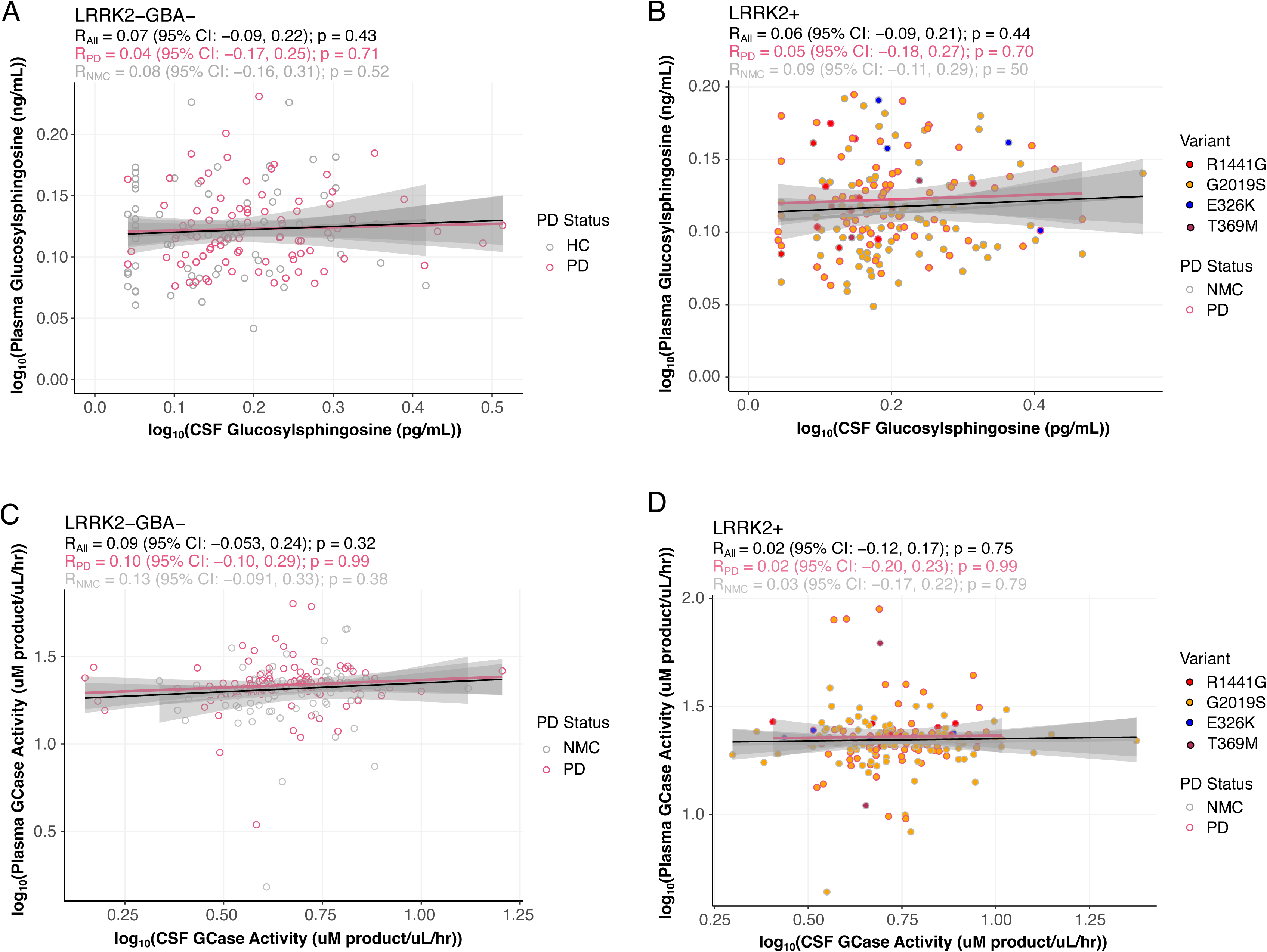
CSF and plasma measurements of GlcSph and GCase activity do not show within-subject correlation in non-carriers and LRRK2 carriers. Correlations between CSF and plasma GlcSph and GCase activity within-subject in non-carrier and LRRK2 carrier groups. Pearson correlations are shown between CSF and plasma GlcSph in **A** non-carriers and **B** LRRK2 carriers. The same is shown between CSF and plasma GCase activity in **C** non-carriers and **D** LRRK2 carriers. Data points are colored by disease status and genetic variant as indicated in the legend. Correlations were assessed on log10-transformed values; correlation coefficients (R), 95% confidence intervals, and Benjamini-Hochberg adjusted p-values are shown within each panel. No significant correlations were observed within-subject across central and peripheral measurements.

**Supplementary Figure 2.**
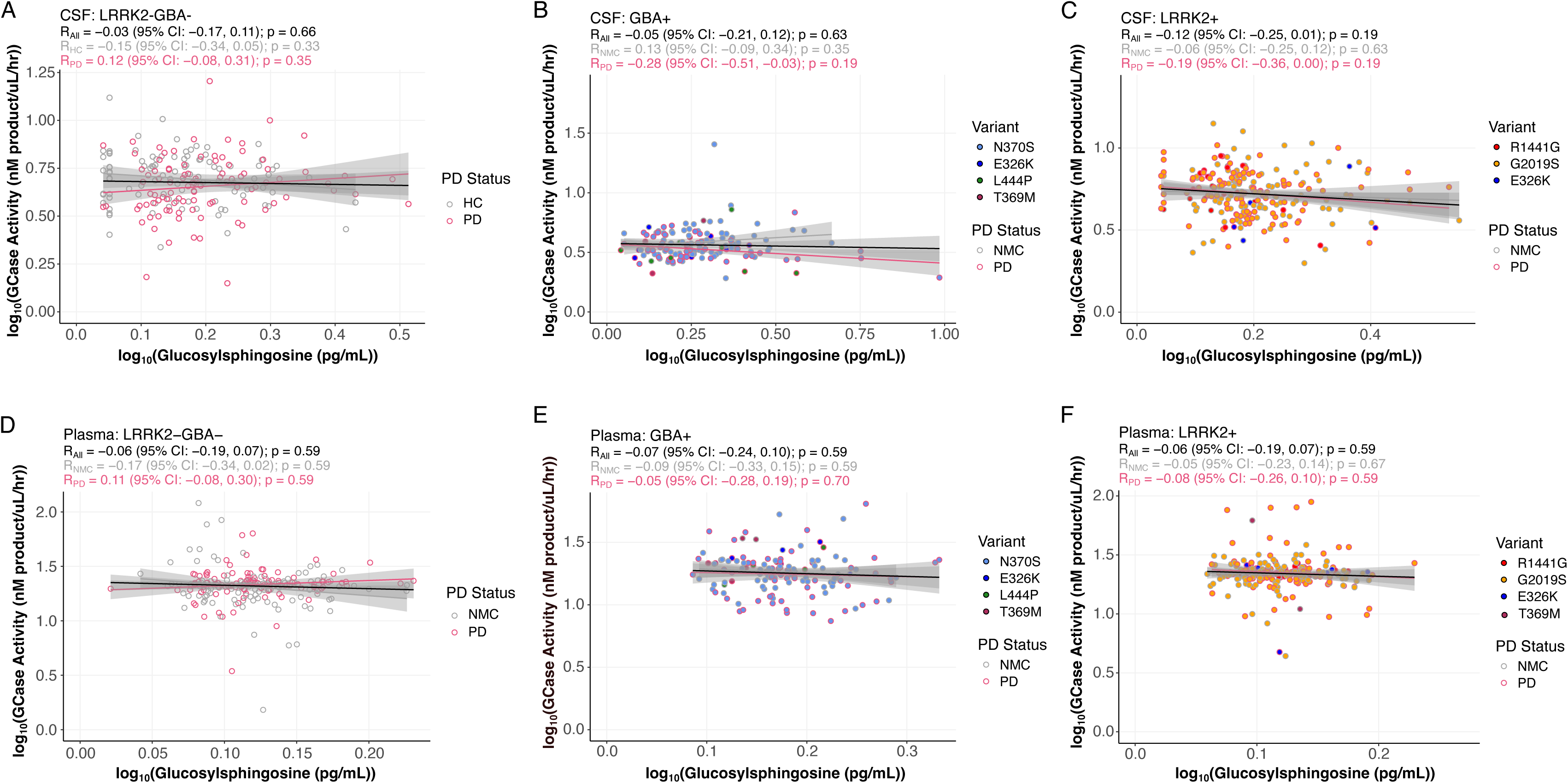
GlcSph levels do not correlate with GCase activity in CSF nor plasma across PD subgroups. Correlations between GlcSph levels and GCase activity in CSF and plasma stratified by genetic subgroup and disease status. CSF correlations are shown for **A** non-carrier, **B** GBA carriers, and **C** LRRK2 carriers. Plasma correlations are shown for **D** non-carrier, **E** GBA carriers, and **F** LRRK2 carriers. Data points are colored by disease status and genetic variant as indicated. Correlations were assessed using Pearson correlation on log10-transformed values; correlation coefficients (R), 95% confidence intervals, and Benjamini-Hochberg adjusted *p*-values are shown within each panel. No significant correlations were observed in any subgroup or biomatrix. Homozygous N370S mutation carriers were omitted from the statistical analyses to prevent correlations driven by this group.

**Supplementary File 1. Summary statistics for sex differences and age correlations within non-carrier and carrier subgroups.** Tab 1 shows ANCOVA pairwise comparisons between sex and adjusted for age. Tab 2 shows the partial Pearson correlations with age.

**Supplementary File 2. Summary statistics for correlation of targeted analytes and clinical assessments.** Partial Pearson correlations were adjusted for age, sex, and PD medication status and Benjamini–Hochberg correction was applied for multiple comparisons. REM_SCORE: REM Sleep Behavior Disorder Screening Questionnaire; MoCA: Montreal Cognitive Assessment; U1_TOTAL: MDS-UPDRS Part 1 total score; MDSUPDRS3_Total: Movement Disorder Society-Unified Parkinson’s Disease Rating Scale Part 3 total score; GDS_Total: Geriatric Depression Scale; QUIP_Total: Questionnaire for Impulsive-Compulsive Disorders in Parkinson’s Disease; U3_FaceSpeech: MDS-UPDRS Part 3 hypomimia and hypokinetic dysarthria score; U3_Rigidity: MDS-UPDRS Part 3 rigidity score; U3_Bradykinesia: MDS-UPDRS bradykinesia score; U3_PostInst: MDS-UPDRS Part 3 postural instability score; U3_Trmr: MDS-UPDRS Part 3 tremor score; UPSIT_rank: University of Pennsylvania Smell Identification Test.

**Supplementary File 3. Summary statistics for 140 PD-associated SNPs against plasma and CSF GlcSph levels and GCase activity.** CHR: chromosome; SNP: rsID; BP: genomic position (hg38); Nearest_Gene: closest gene to highest ranking SNP in the region. A1: effect allele; A2: alternative allele; P_QTL: p-value for SNP to analyte association; BETA_QTL: effect size of SNP to analyte association, in units of SD change of adjusted and transformed analyte levels per dosage of effect allele; P_PD: p-value for PD association; BETA_PD: log odds ratio for PD association; Measurement: analyte, one of plasma GCase activity or GlcSph, or CSF GCase activity or GlcSph.

